# Volumetric two-photon imaging in live cells and embryos via axially gradient excitation

**DOI:** 10.1101/589317

**Authors:** Yufeng Gao, Xianyuan Xia, Jia Yu, Tingai Chen, Zhili Xu, Long Xiao, Liang Wang, Fei Yan, Zhuo Du, Jun Chu, Hairong Zheng, Hui Li, Wei Zheng

## Abstract

Two-photon microscopy(TPM) that features subcellular resolution, intrinsic optical sectioning ability, and deep penetration in sample is a powerful tool of bioimaging. However, the process of layer-by-layer scanning to form a 3D image inherently limits the volumetric imaging speed and significantly increases the phototoxicity. Here we develop a gradient TPM technique that enables rapid volumetric imaging by only acquiring two 2D images. By sequentially exciting the specimen with two axially elongated two-photon beams with complementary gradient intensities, the axial positions of fluorophores can be decoded from the intensity ratio of the paired images. We achieve an axial localization accuracy of 0.728 ± 0.657 μm, which is sufficient for rapid 3D subcellular imaging. We demonstrate the flexibility of the gradient TPM on a variety of sparsely labelled samples, including bead phantoms, mouse brain tissues, live macrophages and live nematode embryos. The results show that, compared with conventional TPM, the 3D imaging speed increases 6 folds while the photobleaching and photodamage are extremely reduced.

## Introduction

Three-dimensional (3D) fluorescence microscopy^1^ has proven essential in biology, allowing interrogation of structure and function at spatial scales spanning macromolecular^2^, cellular^3^, and tissue^4^ levels. Critical factors to consider in 3D microscopy include spatial resolution, determining the smallest feature resolved in the image; temporal resolution, allowing the rapidest 3D information acquisition; and phototoxicity, hindering the applications in live samples. Given inherent tradeoffs in imaging resolution, imaging speed and phototoxicity^5^, no single microscopy technique performs optimally in all these areas. Methods that potentially improve both spatial and temporal resolution while maintaining low phototoxicity to living organisms are thus of great practical interest in biomedical research.

Two-photon excitation fluorescence microscopy (TPM) has been a powerful tool for *in vivo* 3D imaging of cellular and subcellular structures and functions in deep turbid tissues^6^. Attributing to its nonlinear excitation property, TPM provides compelling performance of diffraction-limited spatial resolution. However, conventional TPM captures volumetric images by serially scanning the 3D space with a Gaussian focus, and thus significantly limits the imaging speed which is crucial for capturing rapid biological actions like calcium transient of neurons. A considerable amount of efforts have been made to improve the scanning rate, such as applying resonant scanning, multifocal excitation^7^, and acoustic scanning^8, 9^. Although these methods can partially circumvent the problem by improving the speed of point-by-point scanning that forms a two-dimensional (2D) image, the implementation of layer-by-layer scanning to visualize 3D structures inherently limits the volumetric imaging speed. Moreover, this scanning strategy will repeatedly expose the tissue above or beneath the focal plane to the excitation light and thus significantly aggravate the photodamage and phototoxicity.

Other methods address these limitations by using an axially elongated point spread function (PSF) to capture TPM volumetric image of sparse samples without layer-by-layer scanning^10^. For instance, the Bessel-beam based method^11^ can achieve a video-rate volumetric imaging speed over a 15-400 μm thick volume. However, this methodology can only provide 2D projection images of the samples and lost the information of axial location of fluorophores. Another strategy, volumetric two-photon imaging of neurons using stereoscopy (vTwINS)^12^, used an elongated, V-shaped PSF to encode the depth location of neurons into the separation distance between their 2D projective image pairs. This method can recorded the neuron activity within a 45 μm thick volume at a 30 Hz while preserving the depth location of somas. Unfortunately, to create a V-shaped PSF, the objective NA is partly sacrificed in vTwINS and thus degraded the excitation efficiency and lateral resolution.

Here we present a novel TPM volumetric imaging method based on paired gradient excitation. In contrast to conventional TPM, the axial position of fluorophore is determined by the intensity ratio between paired images rather than layer-by-layer scanning. This strategy increases the volumetric imaging rate at least 6-fold over the conventional TPM and achieves an axial localization accuracy of 0.728 ± 0.657 μm, which is suitable for fast 3D imaging of sparsely labelled samples at subcellular resolution. In addition, our method obviates the out-of-focal plane exposure and therefore significantly decreases the photodamage and phototoxicty caused by repeated exposure. We illustrate the power of our approach by imaging phagocytosis dynamics of macrophages and the development of *C. elegans* embryos.

## Results

The concept of our scheme is straightforward. We first use a phase-only spatial light modulator (SLM) which was conjugated to the back focal plane of the objective to generate a pair of axially elongated focal spots (**Supplementary Fig. 4** and **5**). The intensity of the first focal spot decreases linearly along the optical axis with depth increases, while the second focal spot has an opposite intensity distribution. Then, the specimen was scanned by the paired PSFs successively to obtain two frames of image, which contain depth information in their intensity. Finally, the intensity ratio of the same component in the image pair was calculated and a ratio-depth mapping function was used to determine the axial position of the structure (**Fig. 1a**, **Online Methods**). We refer to this method as gradient TPM (Grad-TPM). In this system, the SLM that served as an additional focus shaping module can be easily assembled into the conventional Gaussian-focus TPM (Gauss-TPM, where 3D image acquisition is based on layer-by-layer scanning) with little lateral resolution degradation (0.76 μm, **Supplementary Fig. 2**). Besides, the switch between Grad-TPM and Gauss-TPM can be fulfilled by conveniently loading specific patterns on the SLM (**Online Methods**).

**Fig. 1.**
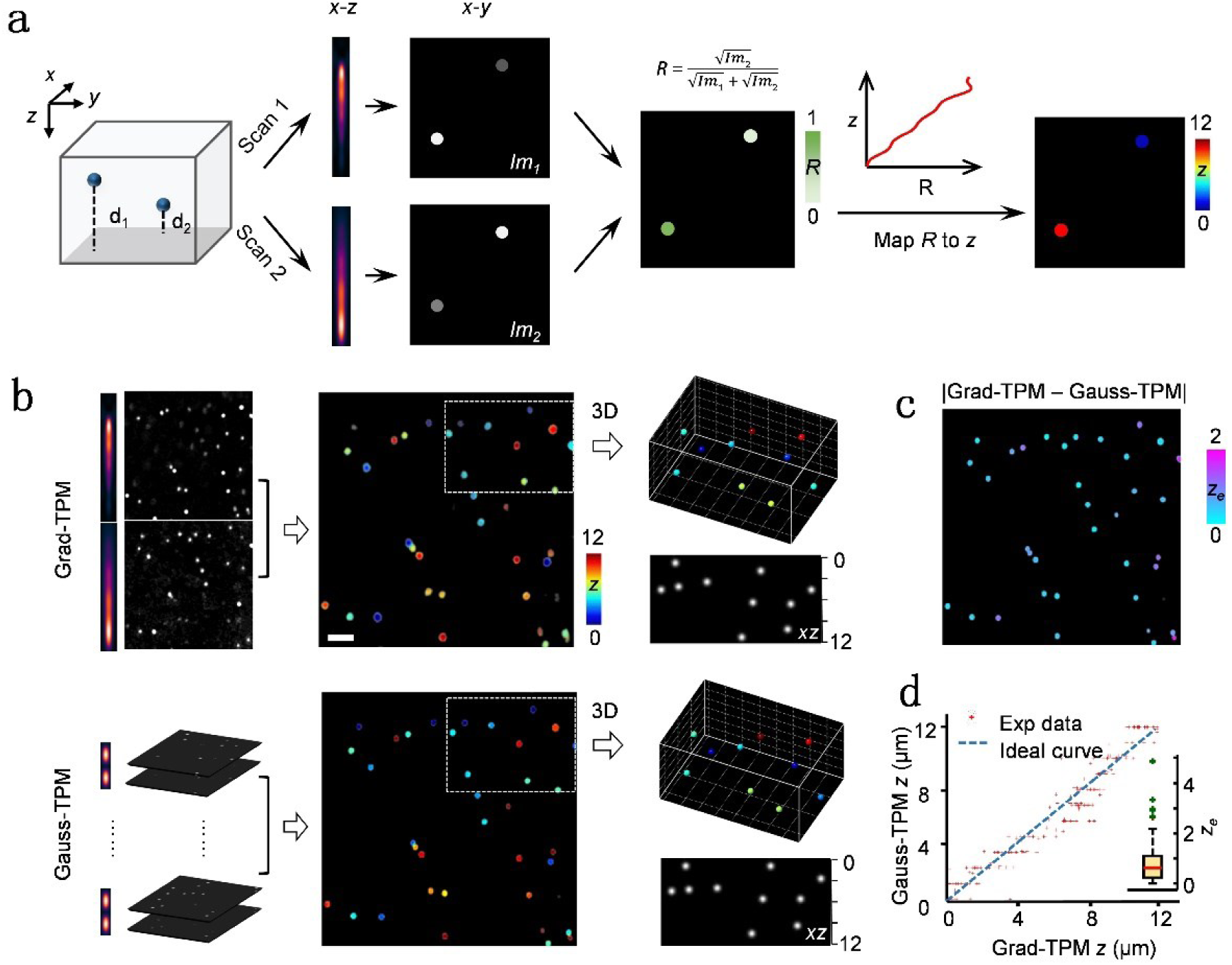
Grad-TPM concept and performance. (**a**) Grad-TPM uses a pair of gradient foci with opposite intensity distributions to successively scan the specimen and thus generates two images. Shallow objects appear brighter in image 1 (*Im*_1_) than in image 2 (*Im*_2_), while the situation is opposite for deep objects. Thus, the depth of a given object can be decoded from the intensity ratio of the two images. (**b**) Fluorescent beads imaged by Grad-TPM (up) and the standard Gauss-TPM (down). (Right panels) The 3D reconstruction and xz view of the boxed region. Scale bar, 5 μm. (**c**) The difference between Grad-TPM depth and Gauss-TPM depth of fluorescent beads (z_e_). (**d**) Gauss-TPM depth as a function of Grad-TPM depth and the boxplot of z_e_. The unit of all the z and z_e_ is μm.

**Fig. 2.**
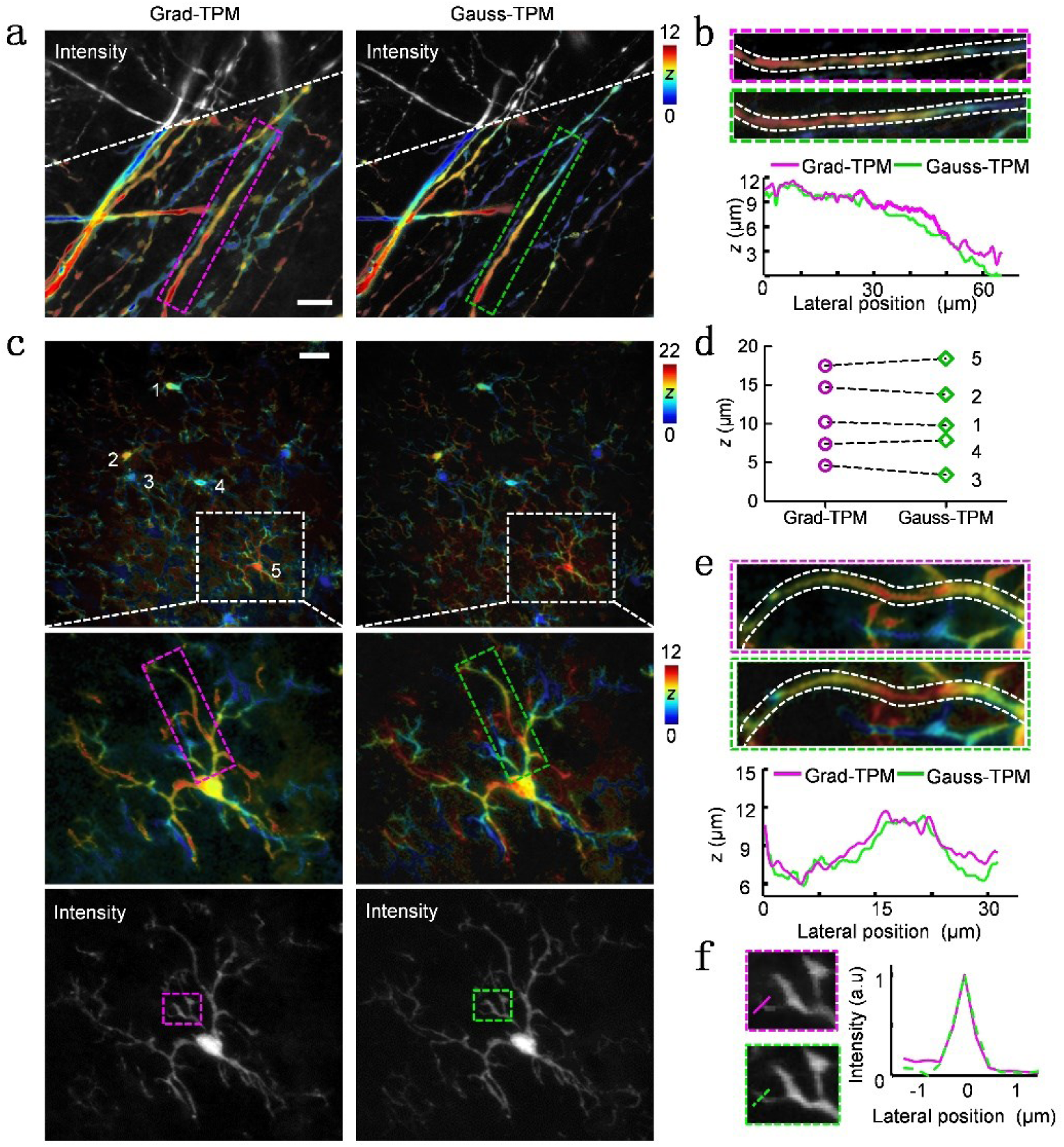
Grad-TPM images of various biological structures show depth resolution and intensity contrast resemble to corresponding Gauss-TPM images. (**a**) Axons in brain slice of Thy1-GFP transgenic mouse. (**b**) Higher magnification views of the boxed axon in **a** (outlined by white dashed lines) and the depth profiles along the central axis of the axon. (**c**) Microglia in brain slice of CX3CR1-GFP transgenic mouse. (**d**) The depths of microglia cell bodies marked by 1-5 in **c**. (e) Higher magnification views of the boxed microglia process in the second row of **c** (outlined by white dashed lines) and the depth profiles along the central axis of the process. (**f**) Higher magnification views of the boxed areas in the third row of **c** and corresponding intensity profiles along the dashed lines for lateral resolution demonstration. Scale bars, 20 μm. The unit of all the *z* is μm.

To verify the feasibility of our approach, we compared the volumetric images taken from Grad-TPM and Gauss-TPM in increasingly complex environments. First, we inspected the performance of our microscope by measuring 1-μm diameter fluorescent beads embedded in the agarose gel (**Fig. 1c-f**). The comparison of the two volumetric images shows that the appearance of each bead in Grad-TPM image is approximately the same to Gauss-TPM result. We further assessed the axial localization accuracy of our approach by comparing the axial center positions of beads from the Grad-TPM image pairs and the reference Gauss-TPM image stacks. The results show that the localization accuracy is 0.728 ± 0.657 μm (372 beads located at depth varying from 0 to 200 μm in the gel, taken from 10 different imaging volumes), which is accurate enough for imaging subcellular structures.

Next, we evaluated the performance of Grad-TPM on various biological samples. Volumetric imaging of neurons and neural networks are crucial to elucidate neural circuit functions. We imaged a fixed Thy1-GFP mouse brain tissue by Grad-TPM and Gauss-TPM, respectively, and compared the 3D architecture of neural networks revealed by these two methods. The fibrous structure of axon extended from the neuron body in three dimensions, and the soma of neuron can be identified clearly. Strict comparison between Grad-TPM image and Gauss-TPM image shows that the derived depth profiles along the axon are almost identical (**Fig. 2a** and **2b**). More than that, in corresponding intensity image, the contrast of different axons is remained without missing any essential or micro structures (**Fig. 2a**).

In addition to fibrous structure, cellular structure was also involved for Grad-TPM performance assessment. The volumetric distribution of microglia in CX3CR1-GFP brain slices were measured (**Fig. 2c-f**). The average depth of each cell body calculated from Grad-TPM images is approximate to the exact depth obtained from Gauss-TPM (the first row of **Fig. 2c** and **2d**) and a precise estimation of the depth variation along the microglia process was also demonstrated (the second row of **Fig. 2c** and **2e**). Note that here, to capture multiple sparsely distributed microglia in the view, we connected two axially adjacent volumes sequentially imaged by Grad-TPM and the results show that the two volumetric images can be stitched seamless in axial direction, which provides the potentiation to form deep 3D image (**Supplementary Fig. 6**).

Having demonstrated the rapid volumetric imaging capability in phantom and fixed biological samples, we next investigated the photobleaching and phototoxicity of Grad-TPM in live cells. Photobleaching and phototoxicity induced by the excitation light is a major problem in fluorescence microscopic imaging of biological samples from cells to organisms, in particular for living systems^3^. Since the proposed Grad-TPM obviated the repeated exposure to out-of-focal plane excitation, we expect that the Grad-TPM may provide much lower photobleaching and phototoxicity relative to the standard 3D imaging method (**Fig. 3a** and **Supplementary Fig. 7**). To validate this assumption, we performed time-lapse fluorescence imaging of live cells with similar signal-to-noise ratio at the beginning. After continually imaging plasmid-transfected HEK293 cells for 25 min, the fluorescence intensities of cells imaged via Grad-TPM slightly diminished by 2.07 ± 2.89 % (n = 7 cells), whereas in the Gauss-TPM 70.85 ± 8.93 % (n = 5 cells) fluorescence was bleached out (**Fig. 3a** and **Supplementary Movie 1**). Even after imaging for 1.5 h, the reduction of fluorescence intensities measured from the Grad-TPM is still negligible (**Fig. 3a**).

**Fig. 3.**
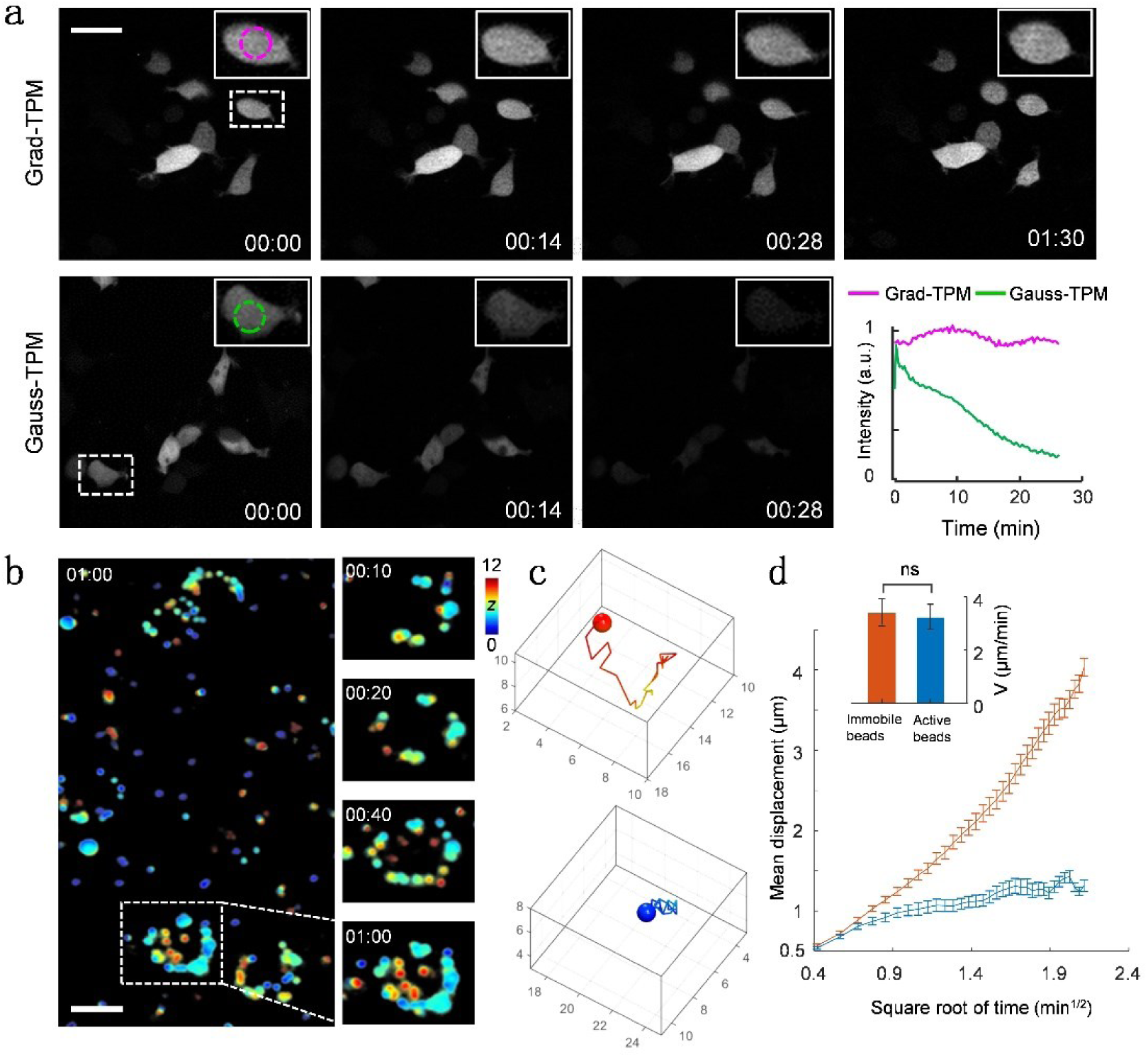
Grad-TPM shows observably lower photobleaching relative to the traditional Gauss-TPM on living cell imaging (a) and thus is highly suitable for longitudinal tracking of biological events, such as the phagocytosis of macrophages (b-d). (**a**) HEK293 cells transfected with pCAG-EGFP DNA recombinant plasmid. The inset on the upper right corner of each image is a magnification view of the boxed area. The circled area in the inset is used to calculate an average intensity to create the line graph. (**b**) Cultured macrophages are phagocytizing fluorescent beads wandering around. Scale bars, 20 μm. The time is shown at the corner as h:min. The unit of *z* is μm. (**c**) Representative trajectories of active beads(up) and immobile bead(down). (**d**) Statistical motility parameters of immobile beads and active beads. Mean velocities indicate that there is no significant difference of bead velocity between these two groups. Mean displacement curves suggest that the immobile beads show constrained motility, while the active beads execute directed migration.

Quantifying the phototoxicity effects on live cells is less straightforward, because the phototoxicity effects depend on many factors such as cell type, imaging conditions, the intracellular distribution and concentrations of fluorophores, and so on^13^. Here, we simply assess the phototoxicity effects by quantifying the rate of death of HepG2 cells during time-lapse imaging. To detect the dead HepG2 cells, propidium iodide (PI), a classical dye that uniquely stains nucleus of dead cells, was applied. To locate the live cells at the start of imaging, we simultaneously labeled the cells with DyLight 488 to highlight their membranes. In Grad-TPM imaging, none of the cells in the view died even at 1.5 h imaging, whereas the cells gradually died after 20 min exposure to Gauss-TPM, suggesting that the phototoxicity is minimal during Grad-TPM imaging **(Supplementary Fig. 7** and **Supplementary Movie 2**). The significantly lower photobleaching and phototoxicity effects of the Grad-TPM on living biological samples make it more powerful and more suitable for obtaining reliable and reproducible quantitative data on biological processes than the standard 3D imaging method.

Benefiting from the rapid volumetric imaging speed and low phototoxicity, the Grad-TPM is highly suitable for longitudinal tracking of embryos development *in vivo*. We performed Grad-TPM imaging on *C. elegans* embryos (strain BV24) over 2 h, with the imaging rate of 0.056volume/s (8 s for scanning and 10 s for interval). Cellular divisions throughout the volume were captured without photobleaching or obvious phototoxicity (**Supplementary Fig. 8** and **Supplementary Movie 3**). We also performed Grad-TPM imaging at a speed of 0.5 volume/s (no interval between each pair of scans) on *C. elegans* embryos for 3.5 h (**Supplementary Movie 4**). There is no sign of obvious photobleaching and phototoxicity during imaging, indicating that the Grad-TPM method is not only adept in tracking prolonged biological processes but also capable of capturing fast or detailed dynamic behaviors. We used the same approach successfully captured the 3D process of macrophages phagocytizing fluorescent beads at a speed of 0.1 volume/s (**Fig. 3b-d** and **Supplementary Movie 5**). The wandering beads gradually gathered together to form a ring and stay immobile afterwards, suggesting that the beads were phagocytized by the nearby microphage and located in its cytoplasm. To quantitatively analyze the bead motilities, we simply classified the beads into two groups, immobile beads and active beads according to the migration radius, which is defined as the straight-line distance of a bead from its starting point to the farthest point it reached during 6 min. The representative trajectories of these two kinds of beads are shown in **Fig. 3c**. By calculating the mean velocity and plotting the mean displacement as a function of square root of time, we found that the immobile beads (migration radius < 6 μm) show constrained motility, while the active beads (migration radius > 6 μm) execute directed migration, although there is no significant difference between the velocity of these two groups^14^ (**Fig. 3d**). These motility features indicate that, the immobile beads might have been “locked” by the macrophages, while the active beads might be being transported into/in the macrophages. The result demonstrates the feasibility of using Grad-TPM to reveal quantitative 3D dynamics and interactions of cells and macromolecules, which is impossible for Bessel-beam based volumetric imaging.

## Discussion

The relative limitation in volumetric imaging speed and phototoxicity of Gauss-TPM considerably offsets its original advantages of 3D imaging of living tissues. Our scheme partially solved this problem by modulating the excitation intensity and integrating the depth information into the fluorescence intensity. This strategy can speed up the volumetric image 6-10 folds relative to Gauss-TPM depending on the focal length (**Supplementary Fig. 3**) while mitigating the photobleaching and phototoxity due to the efficient utilization of excitation photons in the axial direction. In addition, our system can be easily implemented by simply adding a SLM that conjugated to the objective back focal plane in the standard Gauss-TPM. It doesn’t need any other specific design or modification of excitation and emission paths.

In the experiments described throughout this study, the thickness of volumetric image obtained by Grad-TPM is set at 12-μm, which is suitable for the visualization of cultured live cells and multicellular organisms such as embryos. However, this thickness could be easily adjusted from the axial diffraction limit, about 4 μm, to 20 μm by applying different phase patterns on the SLM (**Supplementary Fig. 3**). The maximum value is restricted both by the smallest phase pattern, which the SLM can generate, and the excitation power, where the extended focus will significantly decreases the excitation photon density. In practice, we found the elongated PSF will decrease the accuracy of axial localization and we recommend the modest length of 12 μm which may be excellent for our system setup. For the purpose of imaging deeper volume, stitching in axial direction as demonstrated in **Fig. 2** and **supplementary Fig. 6** is better than elongating the PSF.

We note, however, several limitations or caveats in the current method. First, we map the intensity ratio of structures into their depth position, so the localization accuracy depends on the accuracy of intensity information. Biological samples, especially animal tissue, show complex fluorescence fluctuation which may cause localization errors. And the optical aberrations, caused by system alignment and refraction index mismatch of samples, can further distort the gradient focus. To address this problem, several approaches could be adopted. Adaptive optics^15, 16^ which is used to compensate or reduce the optical distortion would restore the designed PSF in deep tissue. And designing a more sophisticated axial intensity distribution instead of the simple linear distribution used in this study would be helpful to accurately localize the target components (**Supplementary Note)**. Second, the current scheme is suitable for the visualization of sparsely labeled fluorescent samples where only one fluorescent target presented in the axial direction. Otherwise, the axial location information of different fluorescence targets will be confused and undistinguished in fluorescence intensity. It would be an approach to solve this problem by adjusting the PSF length to control the volumetric thickness to ensure the sparsity. However this method will sacrifice the imaging speed. Currently, we are developing a stereoscopic Grad-TPM^12^ which uses two pairs of tilt gradient focus as the excitation source. The depth information is not only integrated in the fluorescence intensity but also involved in the lateral shift patterns. In principle, this strategy can identify dense targets in a single volume. Finally, we should admit that the volumetric imaging speed in our experiment is 0.5 volume/s. It is relatively modest, especially compared with the method that based on an acoustic lens. However, by incorporating with fast scanning strategies such as resonant scanning or spinning disk, the volumetric imaging speed can be improved to more than 30 Hz. In principle, our method can be applied to any layer-by-layer based TPM system to improve the volumetric imaging speed.

In summary, we proposed a new method to capture 3D image by encoding depth information into fluorescence intensity. The method can substantially increase the volumetric imaging speed and decrease the phototoxicity of TPM without resolution degradation. This concept, using intensity ratio to decipher the location information, could be potentially applied to other imaging technologies such as super resolution microscopy for extracting extra location information from conventional image.

## Author Contributions

Conceived project: W.Z, H.L. and Y.G. Designed optical system: Y.G, H.L, T.C, J.Y, Y.W. and W.Z. Built optical system and wrote software: Y.G, T.C, J.Y, and H.L. Acquired data: Y.G and X.X. Analyzed data: Y. G and H.L. Prepared samples: L.X, L.W, and F.Y, Provided advice on biological samples: Z.D, F.Y, and H.Z. Wrote paper: Y.G, H.L and W.Z., with advice from all authors. Supervised research: W.Z.

## Acknowledgements

Support for this work was provided by the National Key Research and Development Program of China (2017YFC0110200); Program 973 (2015CB755502); National Natural Science Foundation of China (81471702, 81571724, 81701744, 81822023); Natural Science Foundation of Guangdong Province (2014A030312006, 2017A030310308); the Scientific Instrument Innovation Team of Chinese Academy of Sciences (GJJSTD20180002); the Shenzhen Basic Research Program (JCYJ20170818164343304, JCYJ20170818155006471); SIAT Innovation Program for Excellent Young Researchers (2016020, 201821).

## Disclaimer

The authors declare no competing financial interests.

## Online Methods

### Optical setup for the Grad-TPM system

The Grad-TPM system was constructed by simply assembling a phase-only SLM (PLUTO-NIR, Holoeye Photonics), which was used for focus shaping, into a standard Gauss-TPM. During imaging, two specific phase patterns calculated from a custom-designed algorithm were loaded sequentially on the SLM to generate a pair of axially gradient foci with opposite intensity distributions. Alternately scan the sample with the pair of foci, paired images containing the axial position information of fluorophores were obtained for reconstructing the volumetric image. The Grad-TPM imaging system can easily switch to Gauss-TPM mode by simply loading a background grating pattern on the SLM, which generates a Gaussian focus. For all time-lapse imaging, a black pattern was loaded on the SLM to block the excitation during imaging interval.

The details of the system are presented in **Supplementary Fig. 1**. In brief, pulsed light from a Ti:Sapphire laser (Chameleon Ultra, Coherent), power controlled by an electro-optic modulator, was expanded to a 1/e^2^ diameter of 30 mm before being reflected at 30° from the normal off the SLM. A half-wave plate was placed before the SLM to orientate the polarization angle of the incident laser accurately matching that of the SLM. After being reflected by the SLM, the excitation light was raster scanned two-dimensionally by a pair of mirror galvanometers (5-mm aperture; TS8203, Sunny Technology). An objective (UAPON 40XW340, Olympus) was then used to focus the scanning light on the specimen for eventual imaging. To insure the phase pattern from the SLM stationary at the objective rear pupil plane, even as the galvanometers scan the light, the SLM, the paired galvanometers, and the objective rear pupil plane were made mutually conjugate by three pairs of relay lens (focal lengths f1 = 150 mm and f2 = 75 mm; f1 = f2 = 85 mm; f1 = 80 mm and f2 = 200 mm). The relay lens were also served as beam expander or reducer to match the aperture of each element. In addition, a field stop located at the intermediate image plane between the first pair of relay lenses was utilized to block undesirable light diffracted and / or reflected from the SLM[1]. In the detection path, the emission fluorescence signals were collected by the same objective and separated from the excitation light by a long-pass dichroic mirror immediately above the objective (T715LP, Chroma). A photon counting photomultiplier (H7421-40, Hamamatsu) was then used to detect the signals after being refocused by an additional lens (f = 75 mm) and spectrally filtered by appropriate bandpass filter (Semrock). Besides, for deep Grad-TPM 3D imaging that connects multiple axially adjacent volumes or Gauss-TPM volumetric imaging based on layer-by-layer scanning, the objective was axially translated by an actuator (KMTS25E, Thorlabs).

### Gradient focus generation

In the Grad-TPM, the gradient focus pair were designed to possess axial intensity distributions which are not only linearly varying but also complementary (**Supplementary Fig. 9**). To engineering the designed gradient focal spot, the phase of incident wavefront was manipulated by the SLM described above. Technically, we modulate the phases of incident beams with different objective convergence angles to generate a focal spot with desirable axial intensity distribution. To facilitate this process, a novel circularly symmetric pupil function was proposed. This pupil function simply divided the objective lens pupil into 40 rings with equal areas and assigning each ring a phase *φ*_*k*_ (*k* = 1, 2… 40). The intensity distribution of the focal spot resulting from the modulated pupil phase *Φ* = {*φ*_*k*_} was calculated according to the Richards–Wolf vector diffraction theory[2]. The problem eventually becomes searching the optimal *Φ* that gives rise to a focus with closest axial intensity distribution to the desirable gradient focus. To achieve this goal, a genetic algorithm (Matlab toolbox provided by Andrew Chipperfield, *et al*.) with the following criterion

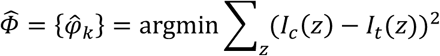

was adopt. Where *I*_*c*_(*z*) and *I*_*t*_(*z*) are the calculated and target axial intensity distributions of the focal spot, respectively. The detailed method is presented in **Supplementary Note** and **Supplementary Fig. 9**. Using the above method, a pair of gradient foci covering 12-μm depth were both obtained in simulation (**Supplementary Fig. 4**) and confirmed in experiment (**Supplementary Fig. 5**).

### Extracting axial location information

To perform Grad-TPM imaging, the specimen was sequentially scanned by the gradient focus pair, *F*_1_ and *F*_2_, and thus generated an image pair, *Im*_1_ and *Im*_2_, which contain depth information in their intensity. More specifically, *F*_1_ possesses intensity that linearly decreases with depth increases, while *F*_2_ has an opposite intensity distribution. In this way, the axial location information is linearly encoded into the intensity of the gradient focus. Since the intensity of two-photon excitation fluorescence is proportional to the square of the excitation intensity, the axial location *z* of a fluorophore is furtherly encoded into its emission intensity and thus can be decoded from the obtained image pair as following:

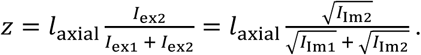

Where *l*_axial_ is the axial length of the gradient PSF, *l*_ex1_ and *l*_ex2_ are intensities of *F*_1_ and *F*_2_ at depth of *z*, respectively, *l*_Im1_ and *l*_Im2_ are the corresponding emission intensities of the fluorophore in *Im*_1_ and *Im*_2_, respectively. Thus, a *z* matrix corresponding to the imaged volume can be obtained.

Since the axial location information is deciphered from the image intensity, the background noise and random noise contained in the image pair would significantly decrease the axial localization accuracy. To attenuate noise interference, three measures were taken (**Supplementary Fig. 10)**. The first one is that only taking area that contains objective fluorophores into calculation. This was done by first normalizing *Im*_1_ and *Im*_2_ to their own maximum intensities, respectively, and then merging them to form a homogeneous excitation image *Im* that reflects the true intensity of fluorophores in the specimen as following (adding up *F*_1_ and *F*_2_ produces a PSF with homogeneous intensity along the optical axis):

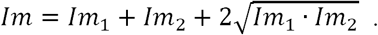

Afterwards, a binary mask of *Im* was created using a threshold and applied to *Im* to eventually isolate objective fluorophores from the background. The second measure is normalizing the calculated *z* in the range of *z*_min_-*z*_max_ to 0-12 μm, where *z*_min_ and *z*_max_ are the minimum and maximum values of *z* matrix that has excluded outliers (please refer to **Supplementary Fig. 10e** as an example). The third measure is that applying a moving average filter with a window size of 3 pixel × 3 pixel to *z* matrix after normalization.

Eventually, to visually present the volumetric imaging result of Grad-TPM, we created a color-coded image which preserves both the spatial location and the intensity contrast information of the specimen by coding depth *z* with colors and intensity *Im* with the saturation of colors, respectively (**Supplementary Fig. 10f**).

### Data acquisition

To assess the performance of Grad-TPM, especially on axial localization accuracy and preservation of intensity and contrast information, Gauss-TPM was considered as a standard 3D imaging method for comparison. For all imaging experiments, 920 nm laser was used for excitation and 450-550 nm bandpass filter was used for spectrally filtering signals before detection, except imaging DyLight 488 labeled membrane of HepG2 cells during phototoxicity assessment. DyLight 488 was excited by 960 nm laser and the emission fluorescence signals were spectrally filtered by a 550-650 nm bandpass filter. It is rather remarkable that, for photobleaching and phototoxicity assessment, the initial fluorescence intensity, pixel dwell time, filed of view, as well as pixel size were particularly kept the same for Grad-TPM and Gauss-TPM imaging. Here, we didn’t use the same excitation laser power on the specimen. Because in such a situation, the density of excitation photons of the gradient focus pair together is obviously lower than that of the Gaussian focus, since the axial length of gradient focus (∼12 μm) is much longer than that of the Gaussian focus (∼1 μm). Other data acquisition parameters for each experiment are provided in detail in **Supplementary Table**.

### Bead motility analysis

Classical motility analysis methods were used to quantify the dynamics of fluorescent beads during studying macrophage phagocytosis. The beads that flashed by were excluded from analysis. We tracked the beads by recording their 3D centroid coordinates in each time-lapse image to obtain their trajectories and calculate the motility parameters[3], such as velocity and displacement. All of the analyses and calculation were carried out using custom MATLAB (MathWorks) programs.

### Preparation of bead samples

To measure the intensity distribution of the gradient focus and evaluate the lateral resolution and axial localization accuracy of Grad-TPM, the fluorescent beads embedded in the agarose gel was used. To prepare the sample, 10 μL suspension of 1-μm or 0.1-μm diameter yellow–green fluorescent beads (1:100 dilution, F8823 or F8803, Invitrogen) were mixed with 1 mL of 1% agarose solution. The mixture was vortexed vigorously and then deposited on a no. 1.5 glass bottom dish (P35G-1.5-14-C, Matek). The gel was allowed to solidify for several minutes and imaged immediately thereafter.

### Preparation of fixed mouse brain slices

To evaluate the performance of Grad-TPM on biological samples, fixed brain slices from Thy1-GFP and CX3CR1-GFP transgenic mice (∼4 week old) were prepared for imaging. The mice were obtained from Professor Yang Zhan and Professor Bo Peng (Shenzhen Institutes of Advanced Technology (SIAT), Chinese Academy of Sciences (CAS), Shenzhen, China), respectively. The mice were maintained in the specific-pathogen-free animal facility of SIAT, CAS. Thy1-GFP mice express EGFP primarily in mossy fibers in internal granule layer of the cerebellum. CX3CR1-GFP mice express EGFP in brain microglia, monocytes, dendritic cells, and NK cells.

To prepare the brain slice, the mouse was first deeply anesthetized with a mixture of 2% α-chloralose and 10% urethane (8 mL/kg) by intraperitoneal injection. Then, transcranial perfusion with PBS and 4% (wt/vol) paraformaldehyde (PFA) in PBS was performed. At the same time, the mouse was sacrificed by this operation. Afterwards, the mouse brain was excised and then fixed with 4% PFA at 4 °C overnight. Finally, 500-μm-thick coronal slices were freehand sectioned by using a brain matrix. The brain slice was then placed on a glass slide and covered with a microscope coverslip for immediate imaging.

All the experiments were performed in compliance with protocols that had been approved by the Guangdong Provincial Animal Care and Use Committee and following the guidelines of the Animal Experimentation Ethics Committee of SIAT, CAS.

### Preparation of live cell samples

We employed three cell lines: HEK293, HepG2, and RAW264. All cells were cultured in high-glucose DMEM (SH30022.01, HyClone) supplemented with 10% FBS (2023-02, Gibco), 50 U/mL penicillin, and 50 μg/mL streptomycin (SH40003.01, HyClone) in an incubator at 37 °C with 5% CO_2_.

For photobleaching evaluation, HEK293 cells were transfected with pCAG-EGFP DNA recombinant plasmid according to the Lipofectamine 2000 protocol (#11668-027, Invitrogen). Here, 200 ng pCAG-EGFP DNA was used per well per glass bottom dish (D35C4-20-1.5-N, In Vitro Scientific). The imaging buffer was phenol red-free DMEM/F12 (Gibco, 11039-021) containing 10% FBS.

For phototoxicity assessment, HepG2 cells with PI labeled nucleus and DyLight 488 labeled membrane were imaged. PI (P21493, Thermo Fisher) is membrane impermeant and generally excluded from viable cells. As such, PI is commonly used for identifying dead cells in a population. DyLight 488 labeled Lycopersicon esculentum (Tomato) lectin (DL-1174, Vector Lab) was used as a counterstaining for live cell membrane. For staining, cells growing to a density of 70-80 % were incubated in the mixture of PI (1:3000 dilution, 500 nM) and DyLight 488 labeled Lycopersicon esculentum (Tomato) lectin (10 μg/mL) for 10 min.

To observe the phagocytosis of macrophages, RAW264 cells mixed with fluorescent beads were imaged. Cells growing to a density of 70-80 % were passaged to two new dishes and continually incubated for 4-5 h. Then, 10 μL suspension of 1-μm diameter yellow–green fluorescent beads (F8823, Invitrogen) were added into the incubation solution and blended with the cells. Finally, after incubating the mixture of cells and beads for 5 min, the sample was immediately imaged.

### Preparation of embryonic nematodes

Nematode strain BV24 *[ltIs44 [pie-1p-mCherry∷PH(PLC1delta1) + unc-119(+)]; zuIs178 [(his-72 1 kb∷HIS-72∷GFP); unc-119(+)] V]* was employed in imaging. The nematodes were raised at 20 °C on nematode growth medium seeded with *E. coli* OP50 in no. 1.5 glass bottom dish (P35G-1.5-14-C, Matek). Before adding nematode embryos, the dish was coated with poly-D-lysine (1 mg/mL in H_2_O, P4408-10MG, Sigma) for ∼10 min to increase adherence to the bottom. For imaging, the nematode embryos were prepared as previously described[4].

### Code availability

Matlab codes for gradient focus generation and axial location information extraction are provided as Supplementary Software. Labview control virtual instruments for the imaging system are available from the corresponding author upon request.

### Data availability

The data that support the findings of this study are available from the corresponding author upon reasonable request.

