## Supplementary Material for "Volumetric two-photon imaging in live cells and embryos via axially gradient excitation"

**Supplementary Fig. 1.** Gradient two-photon excitation microscope setup.
**Supplementary Fig. 2.** Theoretical and experimentally measured lateral resolutions.
**Supplementary Fig. 3.** Simulated gradient foci with different lengths.
**Supplementary Fig. 4.** Simulated gradient focus pair.
**Supplementary Fig. 5.** Experimental gradient focus pair measured using 1 μm diameter fluorescent beads.
**Supplementary Fig. 6.** Axially Concatenate 3D image of microglia using Grad-TPM.
**Supplementary Fig. 7.** Phototoxicity assessment.
**Supplementary Fig. 8.** Monitoring development of *C. elegans* embryos.
**Supplementary Fig. 9.** Gradient focus generation flowchart.
**Supplementary Fig. 10.** Flowchart for axial location information extraction.

**Supplementary Video S1.** Photobleaching assessment.
**Supplementary Video S2.** Phototoxicity assessment.
**Supplementary Video S3.** Longitudinal 3D tracking of nematode embryos development *in vivo* by Grad-TPM at 0.05 Hz.
**Supplementary Video S4.** Longitudinal 3D tracking of nematode embryos development *in vivo* by Grad-TPM at 0.5 Hz.
**Supplementary Video S5.** Long-term 3D monitoring of phagocytosis of macrophages by Grad-TPM.

**Supplementary Table 1.** Data acquisition parameters.

**Supplementary Note .** Gradient focus generation.

**Supplementary Figures:**

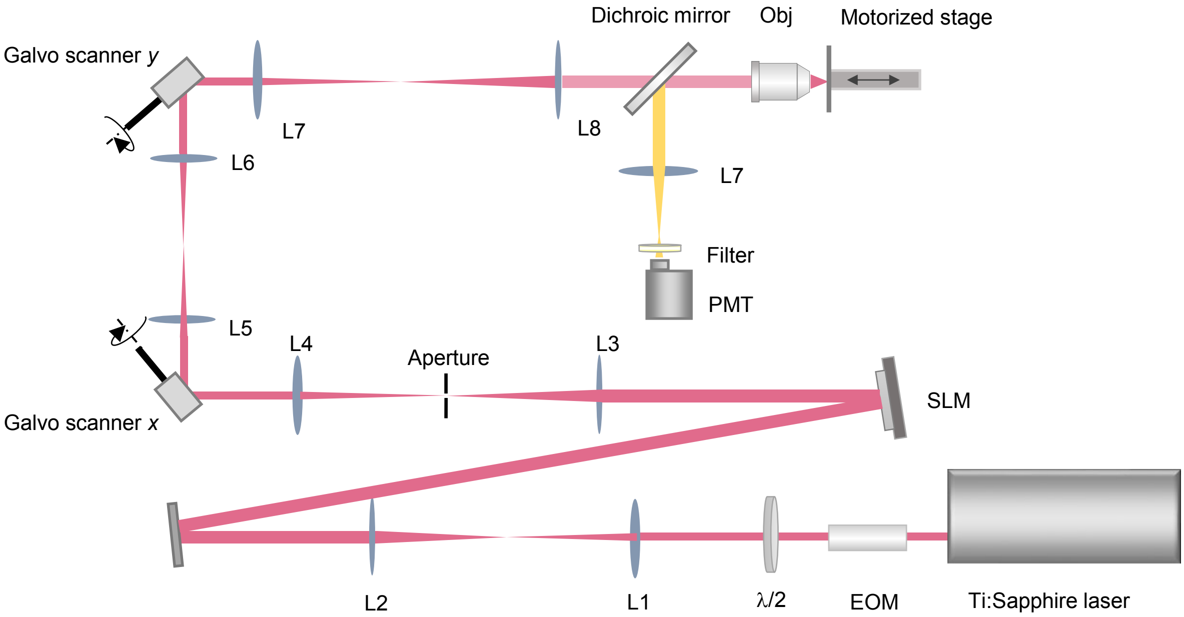


**Supplementary Fig. 1.** Gradient two-photon excitation microscope setup.
EOM, electro-optical modulator; λ/2, 1/2 λ wave plate; L, lens; SLM, spatial light modulator; Obj, objective lens; PMT, photomultiplier.

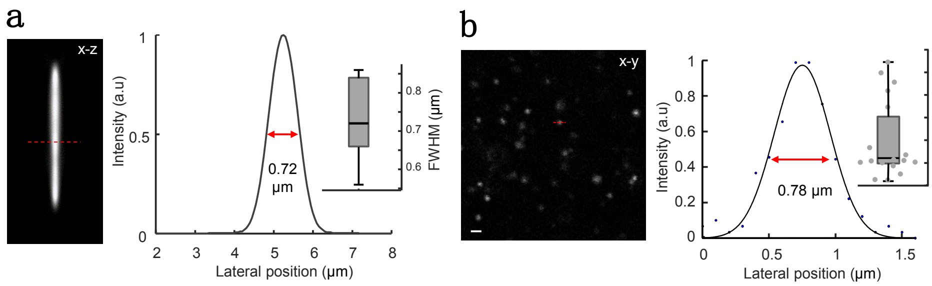


**Supplementary Fig. 2.** Theoretical and experimentally measured lateral resolutions. (**a**) Left, square of the simulated point spread function of Grad-TPM ($PSF^2$). The simulation was based on Richards-Wolf vector diffraction theory and the parameters used in simultion: numerical aperture, 1.0; refractive index of immersion medium, 1.33; excitation wavelength, 920 nm. The operation of squaring is necessary since the intensity of two-photon excitation fluorescence is proportional to the square of the excitation intensity. Scale bar, 1 μm. Right, lateral profile across the middle of the $PSF^2$ (dashed red line on the left panel) and statistical result of lateral resolutions calculated from different depths of the $PSF^2$. (**b**) Left, an exemplary image of 100 nm diameter yellow-green fluorescence beads in gel used for measuring the actual resolution of the Grad-TPM system. Scale bar, 2 μm. Right, profile across the middle of a representative bead (dashed red line on the left panel), corresponding Gaussian fitting result and statistical result of lateral resolution (n = 20).

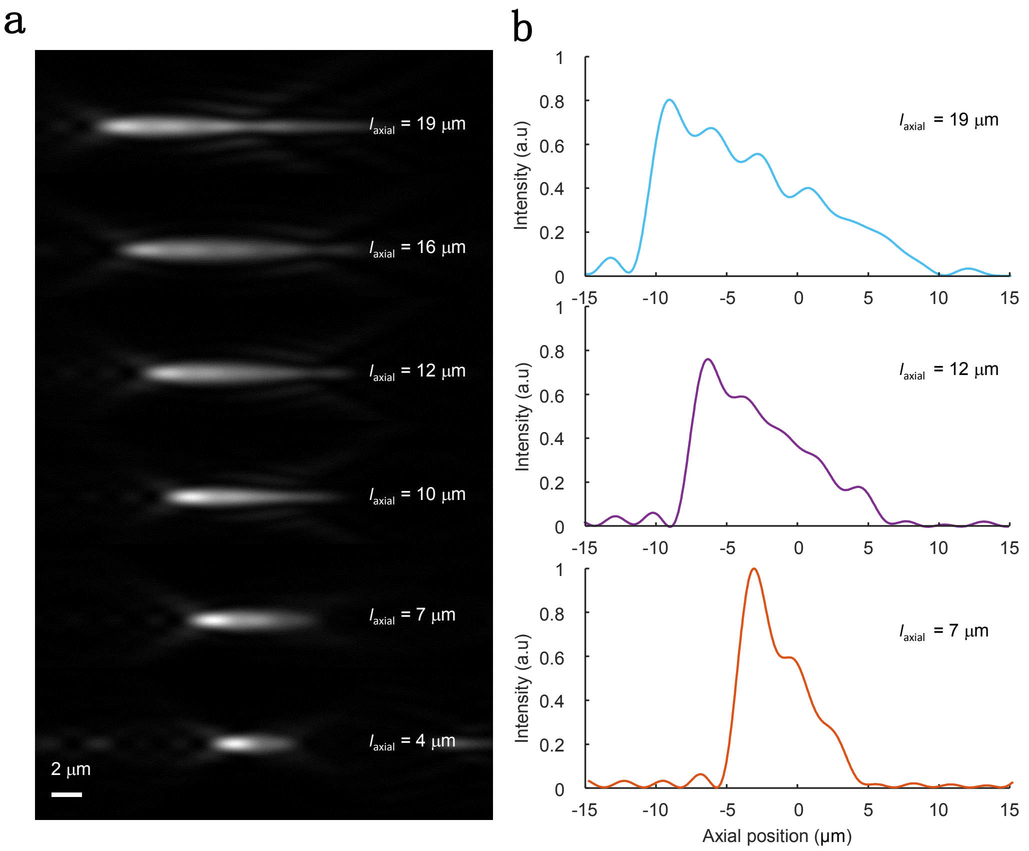


**Supplementary Fig. 3.** Simulated gradient foci with different lengths. (**a**) Intensity distributions that calculated using Richards–Wolf theory. The length of the focus ($l_{\mathrm{axial}}$) can be easily adjusted from 4 μm to 20 μm. Scale bar, 2 μm. (**b**) Corresponding profiles of the 7, 12, and 19 μm gradient foci along the optical axis. Compromising intensity smoothness, photon density and axial length, 12 μm gradient focus is found to be most favorable for our Grad-TPM.

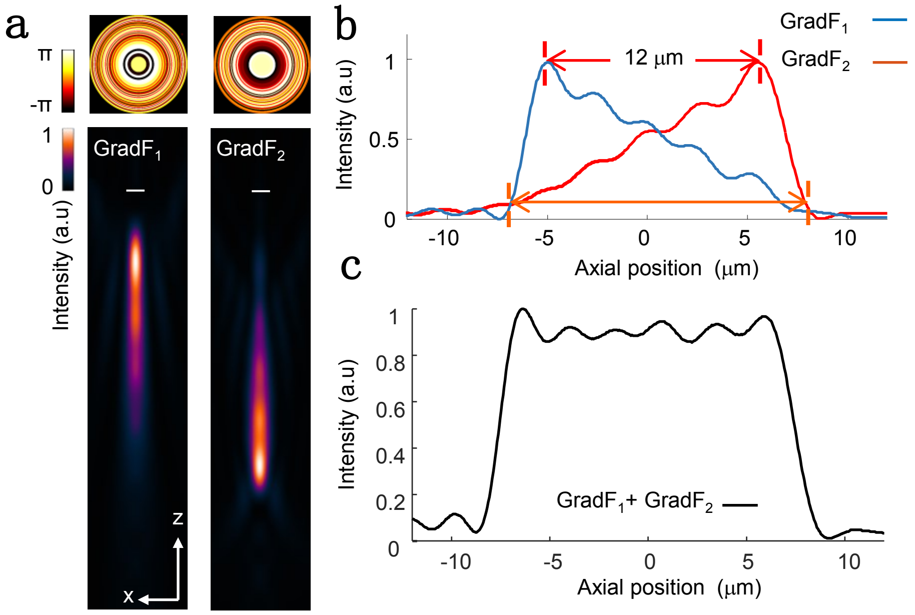


**Supplementary Fig. 4.** Simulated gradient focus pair.
(**a**) Axial views of the PSFs of the pair of gradient foci ($GradF_1$ and $GradF_2$) and corresponding phase patterns that generate the PSFs. Scale bar, 1 μm. (**b**) The intensity profiles of the two gradient foci along the optical axis. (**c**) The sum of the axial intensity profiles of the gradient focus pair.

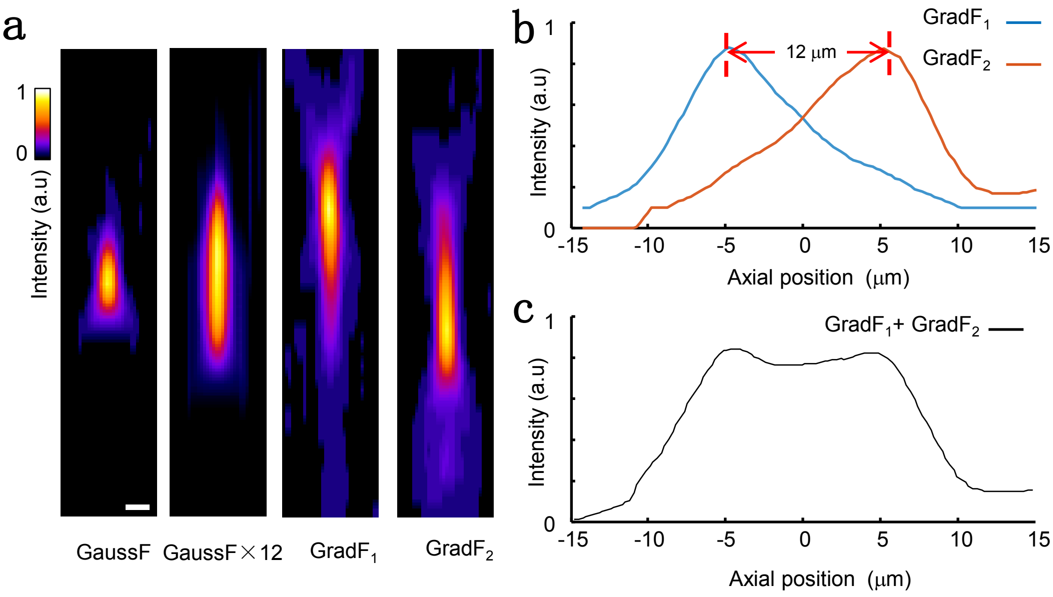


**Supplementary Fig. 5.** Generated actual gradient focus pair measured using 1 μm diameter fluorescent beads.

We measured the PSF of the gradient focus by 3D scanning a 1 μm diameter bead with a voxel size of 0.25 μm × 0.25 μm × 1 μm to make it traverse through the focus. The underlying principle is using the intensity of emission fluorescence to reflect the intensity of the excitation focus. The resulting PSF was then interpolated linearly along the optical axis by a factor of four and filtered by a 3D Gaussian blur with a radius of 1 voxel. Since the intensity of two-photon excitation fluorescence is proportional to the square of the excitation intensity, the square root of obtained images are used to determine the actual distribution of the excitation light around the focus.

(**a**) The axial-view images of 1 μm fluorescent bead indicating the PSF of Gaussian focus (GaussF), axially cascaded Gaussian foci (GaussF × 12), gradient focus 1 and 2 ($GradF_1$ and $GradF_2$) from left to right. Scale bar, 1 μm. (**b**) Measured intensity profiles of $GradF_1$ and $GradF_2$ along the optical axis. (**c**) Measured axial intensity profile of the sum of $GradF_1$ and $GradF_2$.

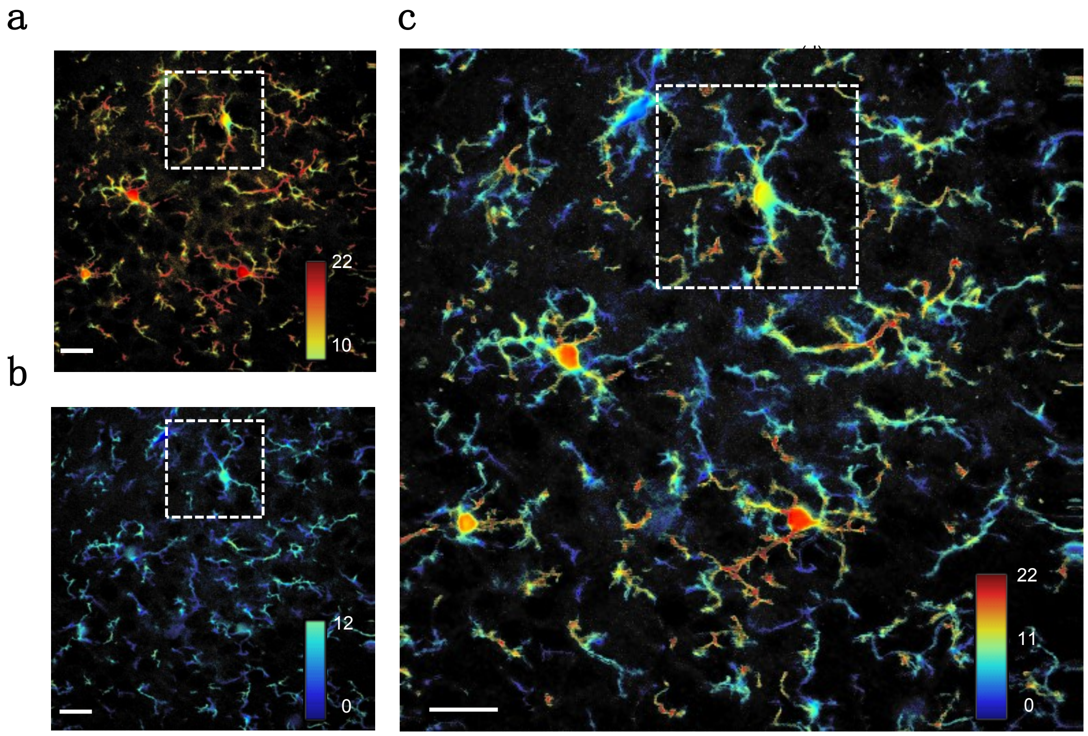


**Supplementary Fig. 6.** Axially Concatenate 3D image of microglia using Grad-TPM. By Axially concatenating contiguous volumes, the imaging view and depth of Grad-TPM can be largely expanded. (**a**, **b**) 3D images of microglia in brain slice of CX3CR1-GFP transgenic mouse covering the depths of 0-12 μm and 10-22 μm, respectively. A 2 μm overlap was particularly set to ensure the junction is intact, smooth and seamless. (**c**) 3D image obtained by concatenating **a** and **b** in axial direction. Scale bars, 20 μm. The unit of all the *z* is μm.

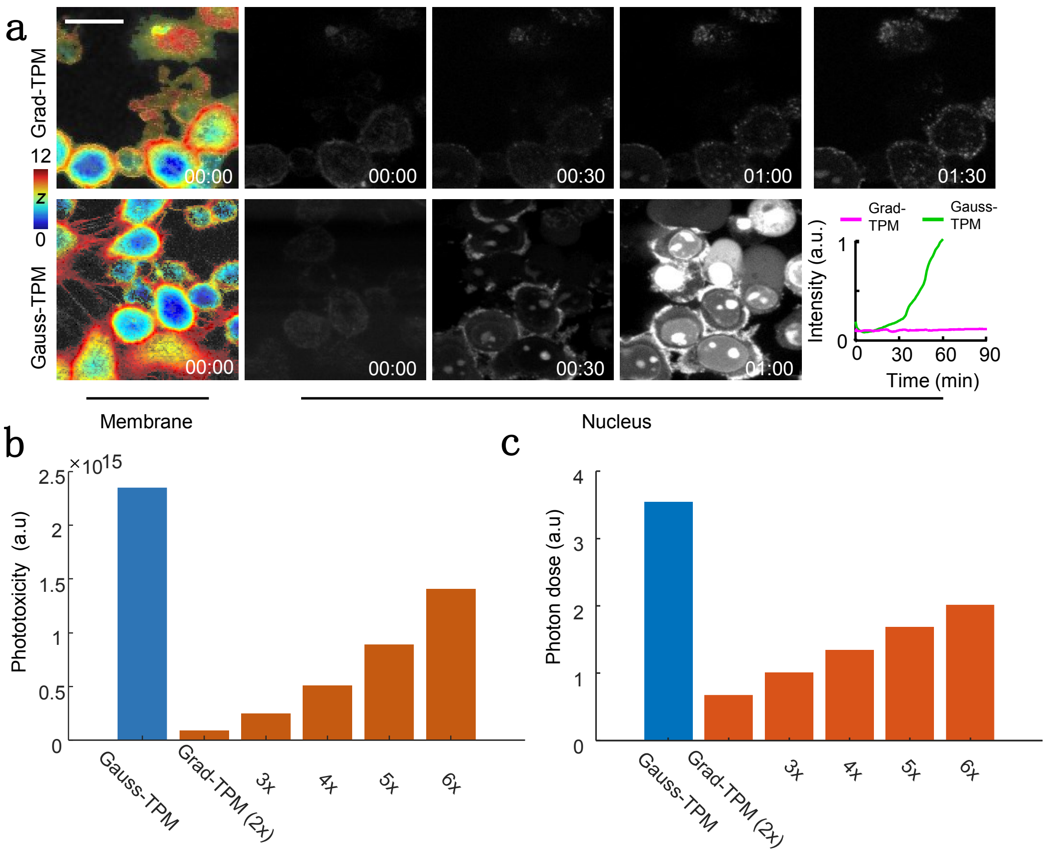


**Supplementary Fig. 7.** Phototoxicity assessment. (**a**) Time-lapse imaging on living HepG2 cells by Grad-TPM shows observably lower phototoxicity relative to the traditional Gauss-TPM. The cells' membrane and nucleus were labeled with DyLight 488 and PI, respectively. PI can't cross the membrane of live cells and appear bright only if it binds to the nucleic acids after the cell dead. The line graph is created by calculating the average intensity of the whole nucleus image at each time point. Scale bars, 20 μm. The time is shown at the corner as h:min. The unit of *z* is μm. (**b**) Theoretical phototoxicity comparison of Grad-TPM and Gauss-TPM. 2-6× indicate the excitation power of Grad-TPM is 2-6 times of that of the Gauss-TPM. (**c**) Theoretical photon-dose comparison. In TPM, phototoxicity has been shown to scale as $\int I(\vec{x},t)^{2.5}\, dV\, dt$, photon dose has been shown to scale as $\int I(\vec{x},t)\, dV\, dt$, where $I(\vec{x},t)$ is the intensity distribution, *V* is the excited volume, and *t* is time[1].

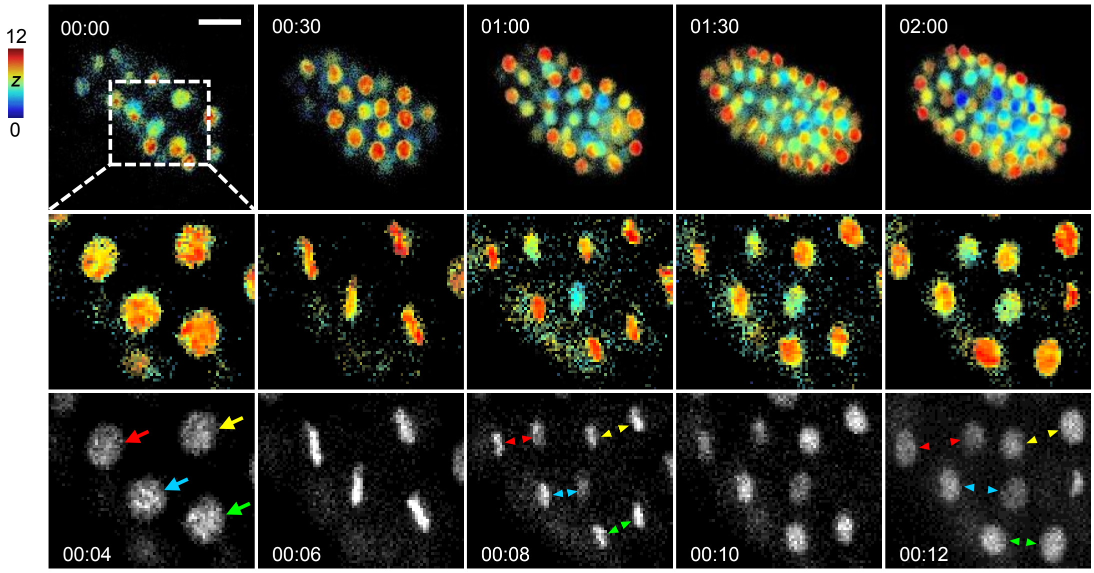


**Supplementary Fig. 8.** Monitoring development of *C. elegans* embryos by Grad-TPM. The development of *C. elegans* embryos recorded lasting 2 h. Mitosis that lasting ~8 min is particularly shown as magnification views at the last two rows. The arrow and arrow head indicate interphase nucleus and condensed mitotic chromatin, respectively. Scale bar, 20 µm. The time is shown at the corner as h:min. The unit of *z* is µm.

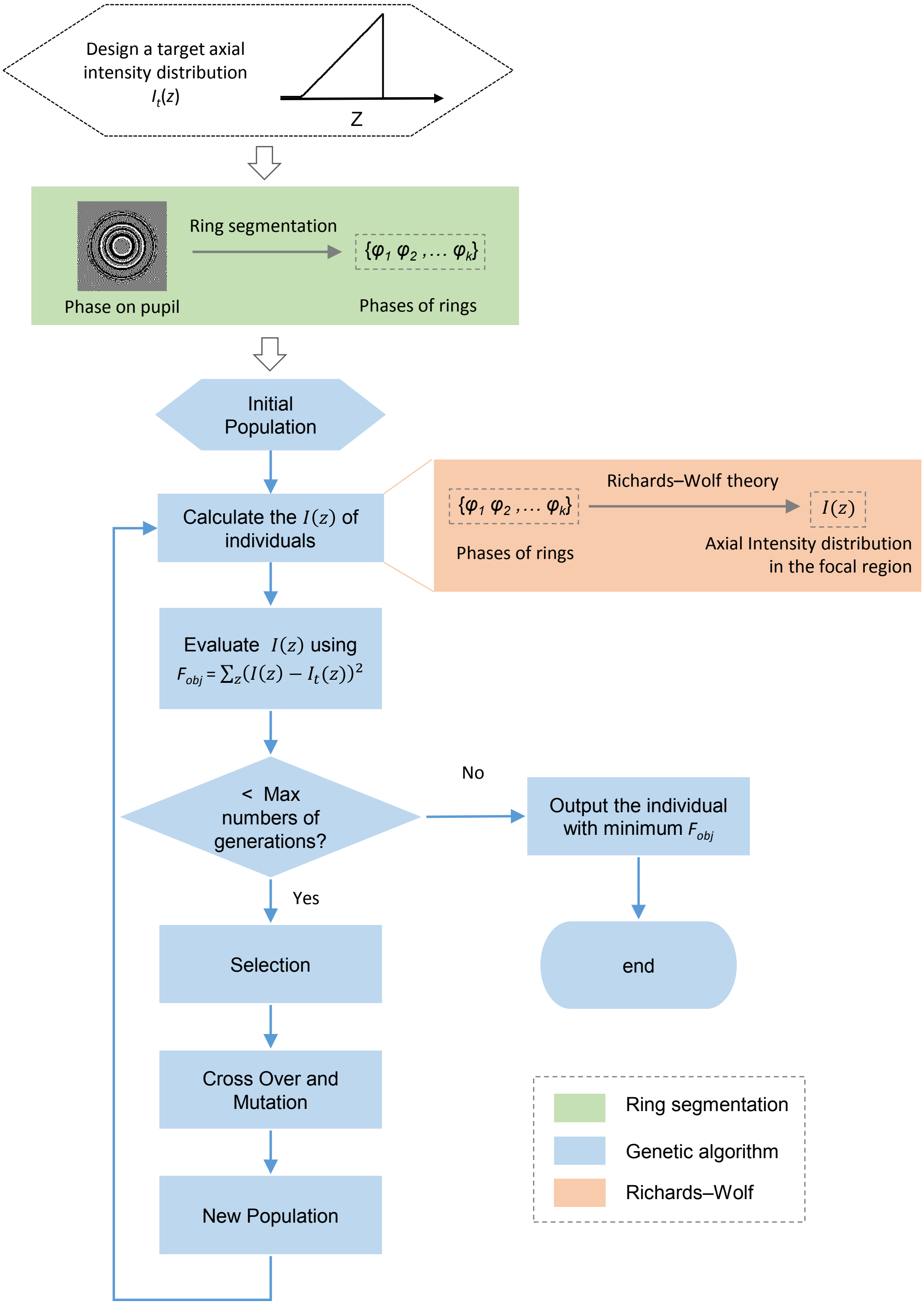


**Supplementary Fig. 9.** Gradient focus generation flowchart (please refer to **Supplementary Note 1** for details).

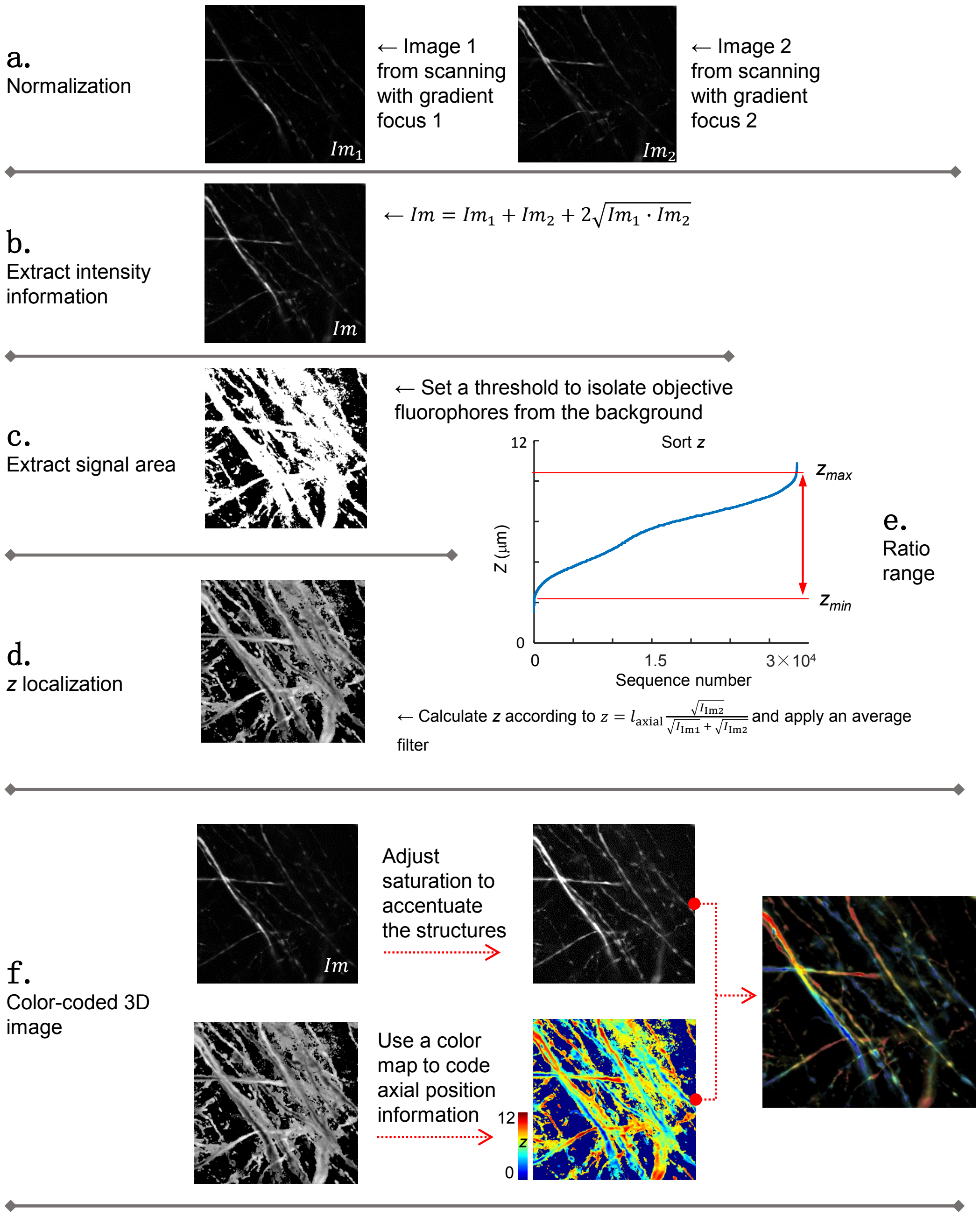


**Supplementary Fig. 10.** Flowchart for axial location information extraction. Color bar unit, μm.

#### Supplementary Videos:

**Supplementary Video S1.** Photobleaching assessment.
EGFP labeled HEK293 cells were imaged by Grad-TPM and Gauss-TPM with the same volume size and acquisition speed. The photobleaching of Grad-TPM imaging is found to be obviously lower than that of the Gauss-TPM imaging and thus more favorable for observing dynamic biological events *in vivo*. The time is shown at the bottom as min:s.

**Supplementary Video S2.** Phototoxicity assessment.
HepG2 cells with DyLight 488 labeled membrane and PI labeled nucleus were imaged by Grad-TPM and Gauss-TPM with the same volume size and acquisition speed. PI can't cross the membrane of live cells and appear bright only if it binds to the nucleic acids after the cell dead. There is no dead cells were observed during Grad-TPM imaging while the number of dead cells gradually increases during Gauss-TPM imaging, indicating that the phototoxicity of Grad-TPM is negligible so that more suitable for long-term monitoring of live systems. The time is shown at the bottom as min:s.

**Supplementary Video S3.** Longitudinal 3D tracking of nematode embryos development *in vivo* by Grad-TPM at 0.05 Hz. The time is shown at the lower left corner as h:min:s.

**Supplementary Video S4.** Longitudinal 3D tracking of nematode embryos development *in vivo* by Grad-TPM at 0.5 Hz. The time is shown at the bottom as h:min:s.

**Supplementary Video S5.** Long-term 3D monitoring of phagocytosis of macrophages by Grad-TPM.
Macrophages are phagocytizing fluorescent beads wandering around. The time is shown at the lower left corner as min:s.

**Supplementary Table 1.** Data acquisition parameters

| **Sample** | **Figure/ Video** | **Imaging mode** | **Volume acquisition speed (s/volume)** | **Interval time (s)** | **Volume size (μm³)** | **Excitation power (mW)** |
|---|---|---|---|---|---|---|
| Bead | Fig. 1c-f | Gauss-TPM | 24 | 0 | 50×50×12 | 2 |
| | | Grad-TPM | 4 | 0 | 50×50×12 | 6 |
| Brain slice of Thy1-GFP mouse | Fig. 2a | Gauss-TPM | 120 | 0 | 200×200×12 | 9 |
| | | Grad-TPM | 20 | 0 | 200×200×12 | 18 |
| Brain slice of $CX_3CR1$-GFP mouse | Fig. 2b | Gauss-TPM | 220 | 0 | 200×200×22 | 12 |
| | | Grad-TPM | ~37 | 0 | 200×200×22 | ~30 |
| HEK293 cell | Fig. 3a/ Video S1 | Gauss-TPM | 13 | 2 | 100×100×13 | ~20 |
| | | Grad-TPM | 2 | 13 | 100×100×12 | 45 |
| RAW264 cell (macrophage) | Fig. 3b/ Video S5 | Grad-TPM | 8 | 2 | 200×200×12 | 6 |
| HepG2 cell | Supplementary Fig. 7a/ Video S2 | Gauss-TPM | 52 | 8 | 62.5×62.5×13 | ~20 |
| | | Grad-TPM | 8 | 52 | 62.5×62.5×12 | 45 |
| Nematode embryos | Supplementary Fig. 8/ Video S3 | Grad-TPM | 8 | ~10 | 150×150×12 | ~40 |
| | Video S4 | Grad-TPM | 2 | 0 | 50×50×12 | ~40 |

### Supplementary Note. Gradient focus generation

Figuring out the pupil phase pattern that generates the gradient focal spot is the crux of the proposed Grad-TPM technique. Conventional methods that based on paraxial theory of scalar diffraction, such as Gerchberg-Saxto phase retrieval [2] and extended Nijboer-Zernike theory [3], perform well in low NA optical systems, but imprecisely in high NA configurations. Here, we developed an algorithm that also works well for high NA systems like the presented Grad-TPM (**Supplementary Fig. 9**). The kernel of the algorithm includes three parts: ring segmentation that facilitates establishing the pupil phase function; Richards–Wolf vector diffraction theory that correlates the intensity distribution of the focal spot and the pupil phase; and the genetic algorithm for searching the optimal pupil phase pattern that gives rise to the desirable gradient focus.

***a. Ring segmentation***

An ideal two-photon microscope can be considered as a rotationally symmetrical system. Therefore, the pupil phase function expressed in polar coordinates can be greatly simplified by only involving the radial coordinate into calculation. Thus, we restate the pupil as a series of area-equivalent concentric rings each having a phase $\varphi_k$ ($k = 1, 2, \ldots$) to form the pupil phase function. The intuition underlying ring segmentation is to manipulate the relative phases of light beams with different objective convergence angles by modulating the pupil phase function, and then recombine the fields they generated to form a predesigned intensity distribution around the focus. Apparently, by increasing the number of rings, we can approach to more and more precise pupil phase functions. 40 rings were adopted in our system, since the minimum width of the rings have to be greater than the size of a single pixel of the SLM.

***b. Richards-Wolf vector diffraction theory***

Richards-Wolf vector diffraction theory [4, 5], a generalization of the Debye integral representation, can precisely describe the electromagnetic field in the image space of a high NA optical system, for which the scalar diffraction theory is deficient. Based on this theory, we built a connection between the pupil phase function and the focal field thereby the focal intensity distribution. To simplify computation, only the axial intensity in the range of 25 m above and beneath the objective focal plane was considered in calculation. We suppose that the illumination is uniform and linearly polarized, and thus the axial intensity distribution of the focal spot can be calculated as $I(\mathrm{z}) = |E(z)|^2$ following the Richards-Wolf theory, where $E(z)$ is the axial field distribution and given by [6]

$$E(z) = \int_0^{\alpha} P(\theta)\,(\cos\theta)^{\frac{1}{2}} \sin\theta\,(1+\cos\theta)\,\mathrm{e}^{-ikz\cos\theta}\mathrm{d}\theta. \qquad (4)$$

$P(\theta) = \varphi_k\ (\theta_{k-1} < \theta < \theta_k)$ is the pupil phase function based on ring segmentation, where $\theta$ is the angle of convergence of the objective lens, namely the aperture angle, $\theta_\mathrm{k}$ is the upper-bound aperture angle of the $k_\mathrm{th}$ ring. $\alpha$ is the maximum $\theta$, which is determined by NA of the objective lens and the medium refractive index, $n$, according to $\alpha = \arcsin\left(\frac{\mathrm{NA}}{n}\right)$. $k = 2\pi/\lambda$ is the wavenumber, where $\lambda$ is the wavelength. In our scheme, NA = 1.0, $n$ = 1.33, $\lambda$ = 920 nm.

***c. Genetic algorithm***

Genetic algorithms (GAs) are stochastic global search and optimization methods that mimic the metaphor of natural biological evolution [7]. GAs are well suited for large-scale optimization problems and thus attractive to optimize phases in the focusing task [8]. In this study, we adopted a GA provided by Andrew Chipperfield *et al.* [9] to determine the pupil phases of the 40 subregions divided by the ring segmentation.

As shown in **Supplementary Fig. 9**, the GA in our case began with generating an initial population that contains 8000 individuals. Each individual is a pupil phase function and has 40 chromosomes, which correspond to phases of the 40 rings. Then, $I(z)$ of each individual were calculated according to the Richards-Wolf theory and their similarity to the target profile $I_t(z)$ were measured through an objective function $F_{obj} = \sum_z (I(z) - I_t(z))^2$ to assess the fitness of these individuals for the following selection. Here, both $I(z)$ and $I_t(z)$ were normalized to their maximums. Next, the GA generated a new population for the next round of iteration through selection, crossover and mutation, all of them were performed via default functions provided by the toolbox. The size of the population reduced to 400 when the number of generation reaches 10 and furtherly reduced to 50 when it reaches 50. The large initial population size avoids convergence to a local maximum and the latterly reduced size speeds up the algorithm. Eventually, the iteration stopped when the number of generation reaches 4000 and the individual with minimum $F_{obj}$ in the last population was figured out as a global best satisfactory.

The gradient focus generation algorithm can converge to a globally optimized pupil phase function regardless of the randomly generated initial population at the cost of being time-consuming. Nevertheless, the imaging speed will not be restricted, since the processes of pupil phase optimization and Grad-TPM imaging are totally independent.
